# How social and economic policies have affected the genome of mezcal agaves: the contrasting stories of Bacanora and Espadín

**DOI:** 10.1101/2025.04.30.651404

**Authors:** Anastasia Klimova, Jesús N. Gutiérrez Rivera, Erika Aguirre-Planter, Luis E. Eguiarte

## Abstract

Plant domestication in Mesoamerica gave the world crops of global significance, such as maize, beans, squashes, cocoa, and cotton. Additionally, it has introduced species of regional economic importance, which still display intermediate levels of domestication, including Agave, Opuntia, columnar cacti, Amaranthus, and various ornamental species. Agaves, in particular, hold immense cultural and economic significance in Mexico and play a crucial ecological role in wild plant communities. However, current agricultural practices have negatively impacted both wild populations of agave through overexploitation and habitat destruction, as well as cultivated plants by reducing available landraces and promoting the use of homogeneous plant material. Using genomic data (RADseq) and over 50,000 SNPs, we aimed to assess how local social and political decisions may have influenced genomic diversity and differentiation in intensively managed, clonally propagated *Agave angustifolia* (Espadín) in the state of Oaxaca, as well as in the cultivated *A. angustifolia* used to produce mezcal known as Bacanora in the state of Sonora and their wild counterparts from both regions. We found evidence suggesting that Espadín recently aroused through farmer selection of a clonal lineage with desirable mezcal production attributes (i.e., hybrid vigor), apparently from a cross between genetically distinct wild populations or by hybridization between wild and cultivated varieties. Espadín samples were represented by closely related heterozygous genotypes, with considerable genetic differentiation from wild plants. On the other hand, the genomic composition of cultivated Bacanora agave apparently was influenced by a recently lifted ban (in 1992) on its cultivation and distillation, which, along with the relatively lower popularity of this beverage, allowed cultivated Bacanora agave to maintain the genetic diversity found in wild populations of the regions. We found that social and political decisions may have important impacts on crop genomic diversity and differentiation.

## Introduction

The domestication of plants and animals has been considered the most important achievement in human history over the last 13,000 years [1]. Plant domestication is the human-driven selection process that transforms wild plants into domesticated crops by modifying their morphological, physiological, and genetic traits [2,3]. Domestication is usually a gradual process, which can be characterized by stages consisting of (a) wild harvesting; (b) conscious and unconscious selection to modify plant characteristics (e.g., flowering time, dormancy, size, and changes in defensive structures); and (c) conscious selection of plant material and genetic alteration, where the plant generally loses its ability to survive without human care [4]. Domestication processes at these stages are still ongoing, encompassing a wide range of plant species and agro-biological systems that range from traditional to large-scale industrialized agriculture.

Plant domestication resulted in the independent invention of agriculture by diverse cultures in different parts of the world [1]. Throughout history, plant domestication was originally focused on traits desired by early farming communities [5]. These traits are known as the “domestication syndrome”, which can help us to differentiate domesticated plants from their wild ancestors [6]. The most common domesticated traits in plants are the loss of seed dormancy, increased organ size, decreased seed dispersal, uniformity in growth, change in photosensitivity, and loss of defense mechanisms [5,7]. The domestication process and the morphological and physiological changes that domesticated species have undergone are reflected in their genomes [8,9].

The most common genetic consequence of domestication is a genetic bottleneck, in which genetic variation in the cultivated populations is reduced [10]. This bottleneck is mainly a consequence of a small number of wild progenitors used during domestication and a low effective population size maintained for several generations, which reduces the available gene pool [11,12]. However, the extent to which genetic diversity is lost varies among species and can be mitigated by allowing gene flow between wild and cultivated individuals. On the other hand, additional bottlenecks may occur when crops are transferred from one region to another or bred for specific uses (e.g., when specialized lines are created). In addition to the neutral effects of genetic drift, artificial selection also decreases genetic diversity in domesticated plants by removing variation in traits undesirable for cultivation [13,14].

Mesoamerica is one of the areas of the New World where agriculture was first practiced and one of the main centers of plant domestication [15,16]. This fact was attributed to the region’s large variety of cultures and high biological diversity [17]. Mesoamerican people domesticated more than 200 plant species and currently use more than 5,000 plant species [3,17,18]. Besides crops of global importance, such as maize, beans, chili peppers, squashes, cocoa, and cotton, in Mesoamerica, there are also species of regional economic importance displaying intermediate to advanced levels of domestication, including *Agave, Opuntia, Leucaena*, columnar cacti, Amaranthus, and numerous ornamental species [17].

Among the species domesticated or in the process of domestication in Mesoamerica, the genus *Agave* stands out due to its cultural and economic importance and its essential ecological role in the wild. The rich history of the interaction between humans and agaves dates back thousands of years [19, 20]. All *Agave* species are native to the American continent, and Mexico is this family’s center of origin and adaptive radiation [19,21]. *Agave* is a species-rich genus [22,23,24] that was fundamental for the survival of the original inhabitants of North and Central America [19,25]. For thousands of years, agaves have played a central role as a multipurpose plant that provided building material, food, medicine, decoration, fibers, and beverages to Mesoamerican cultures. For example, *Agave* fiber production has a long history that can be traced from ancestral Pueblo people to modern indigenous groups and geographically from southern to northern Mexico [25,26]. For instance, henequen was already produced from *A. fourcroydes* during the Mayan era, approximately 5,000 years ago [26,27,28]. Today, in Oaxaca state alone, at least 19 species of agave are used for food preparation, four species are used to make fermented beverages, and 16 species are used to make distilled alcoholic beverages [29,30].

Among the products made from *Agave*, alcoholic beverages, in particular distilled spirits, have the most important social, economic, and ecological role, with a great, but sometimes intricated, commercial and geographical history [31,32,33]. The generic term for these distilled drinks is “mezcal”, which derives from Nahuatl and means cooked or baked maguey [34,35]. Mezcal is an alcoholic beverage obtained from the distillation of the stems of different species of *Agave*, previously cooked and crushed or ground [36]. This general definition covers all distilled beverages obtained from agaves, such as tequila, raicilla, bacanora, and mezcal [35,37].

The production of distilled spirits from agave was apparently common in Central Mexico in the 17th and 18th centuries [28,38]. However, soon after, agave distillates were banned due to competition with grape-based spirits imported from Spain. As a result, agave alcoholic beverages were manufactured clandestinely in remote regions, away from the influence of colonial authorities [39,40,41,42]. From the mid-19th century and throughout the 20th century, the legalization of the production of agave distillates changed in independent Mexico, and its growing national and international demand (particularly tequila, mainly made in the Tequila area in Jalisco) became one of the most important economic activities in western Mexico [38,43]. In the mezcal industry, at least 40 species of agaves are now used [41]. However, due to specific ecological and biological characteristics, some species have been more important and used than others [35,43]. Thus, *Agave angustifolia* stands out as one of Mexico’s most important agave species [44].

*Agave angustifolia* is the presumed ancestor of the most widely used mezcal and tequila agave species and varieties. It has the broadest geographic distribution within the genus. This species has great ecological versatility, and it is found from Sonora and Tamaulipas, in the North of Mexico to Costa Rica [19], growing at elevations ranging from sea level to 2,200 m, associated with a variety of vegetation types, including coastal dunes, thorn scrub, subtropical scrub, tropical deciduous forest, and in the transition of these to the *Quercus*-*Pinus* forest [19].

*Agave angustifolia* has a rich ethnobotanical history and a variety of uses [45,46], and it is considered the ancestor of *A. tequilana* var. *azul*, the plant used for tequila production in Jalisco and other areas. *Agave angustifolia* is also the most important species used to produce mezcal in the southern state of Oaxaca, the leading mezcal producer that accounted for more than 90.5% of production in 2023 [44], in particular using a cultivar apparently derived from *A. angustifolia* known as “Espadín” [35, 44].

Although mezcal is considered an ancestral drink, the mezcal of Oaxaca--as we know it today--is a relatively recent beverage developed at the end of the 20th century [47]. From 1950 to 1980, mezcal production in Oaxaca was done in small family factories [48] that produced an inexpensive, folklorized version of tequila [47]. The modernization of tradition took place after the crisis that affected the mezcal sector between 1980 and 1990. Production was under pressure from industrialization, and rising prices were caused by the local shortage of agave plants induced by their export to Jalisco to make tequila. The mezcal sector was modernized, and paradoxically, “traditional mezcal” was invented [47,49].

Modernization and adherence to the structural standards of Designation of Origin of Mezcal (Denominación de Origen de Mezcal, DOM) and Oficial Mexican Norm (Norma Oficial Mexicana, NOM) have allowed mezcal to enter the international market, where it has been a success. Thus, registered production increased from 980 thousand liters in 2011 to 12,239 thousand liters in 2023, with over 90% of mezcal produced in Oaxaca, from which over 86% of the mezcal is distilled from *A. angustifolia* [44].

In Sonora state, *A. angustifolia* is the only species used to produce Bacanora, a lesser-known mezcal but now very popular in northwestern Mexico. The production of Sonoran liquors began around 1800 [50]. The Bacanora industry had a great boost in the decade between 1900 and 1910, with at least 75 distilleries generating significant income. The increase in production apparently also implied the development of agave plantations since wild plants could no longer supply the growing industry. However, the number of hectares dedicated to cultivation at those times is not known [31,51]. In 1915, the Bacanora industry took an unexpected turn when the governor of Sonora, Plutarco Elías Calles, promulgated a “Prohibition Law”, suggesting that alcoholic beverages were the cause of a moral decline. In 1919, Governor Adolfo de la Huerta repealed the decree and allowed, although in a restricted way, the production and trade of beer, table wines, cider, and champagne, except for mezcals and other high-alcohol liquors, including Bacanora [31,51]. Distilleries and agave plantations were destroyed, and producers were persecuted, imprisoned, and sometimes even executed. Prohibition did not extinguish Bacanora, which was produced and sold, for the most part, in a very local manner in inaccessible mountain regions.

After over 75 years of prohibition, the Alcohol Law of 1992 formally ended the ban on Sonoran mezcal [31], and then, in 2000, the Bacanora Designation of Origin was issued [52]. The DO grants the exclusive right to produce Bacanora to 35 Sonora municipalities, covering 38% of the state’s territory. The history of Bacanora in Sonora has been highly dependent on social policy and people in charge of laws, with some favoring production and increasing distillation and the number of plantations. In contrast, other laws pushed production underground, relying only on wild plants. Currently, the production of Bacanora is growing, and while some producers are adopting monoculture clonal reproduction of their crop, similar to that employed in the tequila industry, others still rely on wild plants and seeds and readily include them in cultivation.

Although Sonora and Oaxaca states have had contrasting mezcal histories, both are experiencing an increase in production, particularly Oaxaca, with exponential production growth [44]. This growth is known as the “mezcal boom”, which led both states to an overexploitation of natural resources [53,54,55]. The growing demand for mezcal has increased the overexploitation of wild agaves and their seeds through overcollection and increased firewood extraction, promoting deforestation. This phenomenon exactly reproduces the expansion of tequila, which promoted the destruction of thousands of hectares of natural vegetation to convert them into agave monocultures. The overexploitation of agaves for tequila production basically extinguished the local wild populations, so now, agave production for tequila is being carried out using clones, which reduces genetic diversity, affecting the capacity of plants to adapt to environmental changes and the attack of pathogens [33,56,57]. Over time, depletion of soil fertility and pest attacks encouraged reliance on agrochemicals (fertilizers, insecticides, and herbicides), contaminating the soil and water and negatively impacting pollinators [58].

One of the main goals of studying plant genetic resources is to understand where, when, and how plants were domesticated from their wild ancestors and how domestication and modification for modern agriculture have affected their genomes. We hypothesized that the contrasting history of cultivation of *A. angustifolia* and mezcal distillation in Sonora and Oaxaca may have been reflected in the genome of the local plants. Our study evaluated how management and social history differences have affected the genetic diversity and differentiation between cultivated and wild *A. angustifolia* plants from Sonora and Oaxaca states. To achieve this, we used more than 50,000 SNP-type genetic markers genotyped in over 200 wild and cultivated *A. angustifolia*. Using a panel of state-of-the-art bioinformatic tools, we found contrasting levels of genetic diversity and differentiation for the cultivated plants within each state. While Espadín presented a clear sign of domestication bottleneck and strong differentiation from the region’s wild plants, Bacanora agaves were found to be genetically diverse but harboring only a portion of the large genetic diversity found in Sonora’s wild plants. Our findings have important implications for the future agricultural management of agaves. They highlighted the urgent need to conserve wild populations of *A. angustifolia* and emphasized the need to develop a breeding program to mitigate the loss of genetic diversity in cultivated plants.

## Materials and methods

### Sample data, SNPs genotyping and filtering

Raw reads from 207 wild and cultivated *A. angustifolia* individuals from the Sonora and Oaxaca states published by [59,60] were downloaded from the Sequence Read Archive (SRX20715033 and SRX20274545). The wild samples from Sonora included 19 sampling sites and 56 individuals. The wild *A. angustifolia* from Oaxaca included 19 sampling sites and 62 individuals (Fig 1). The cultivated samples of *A. angustifolia* included 42 plants from 13 plantations in the Sonora state. The cultivated Espadín samples included 47 plants from 15 plantations in Oaxaca state.

**Fig 1.**
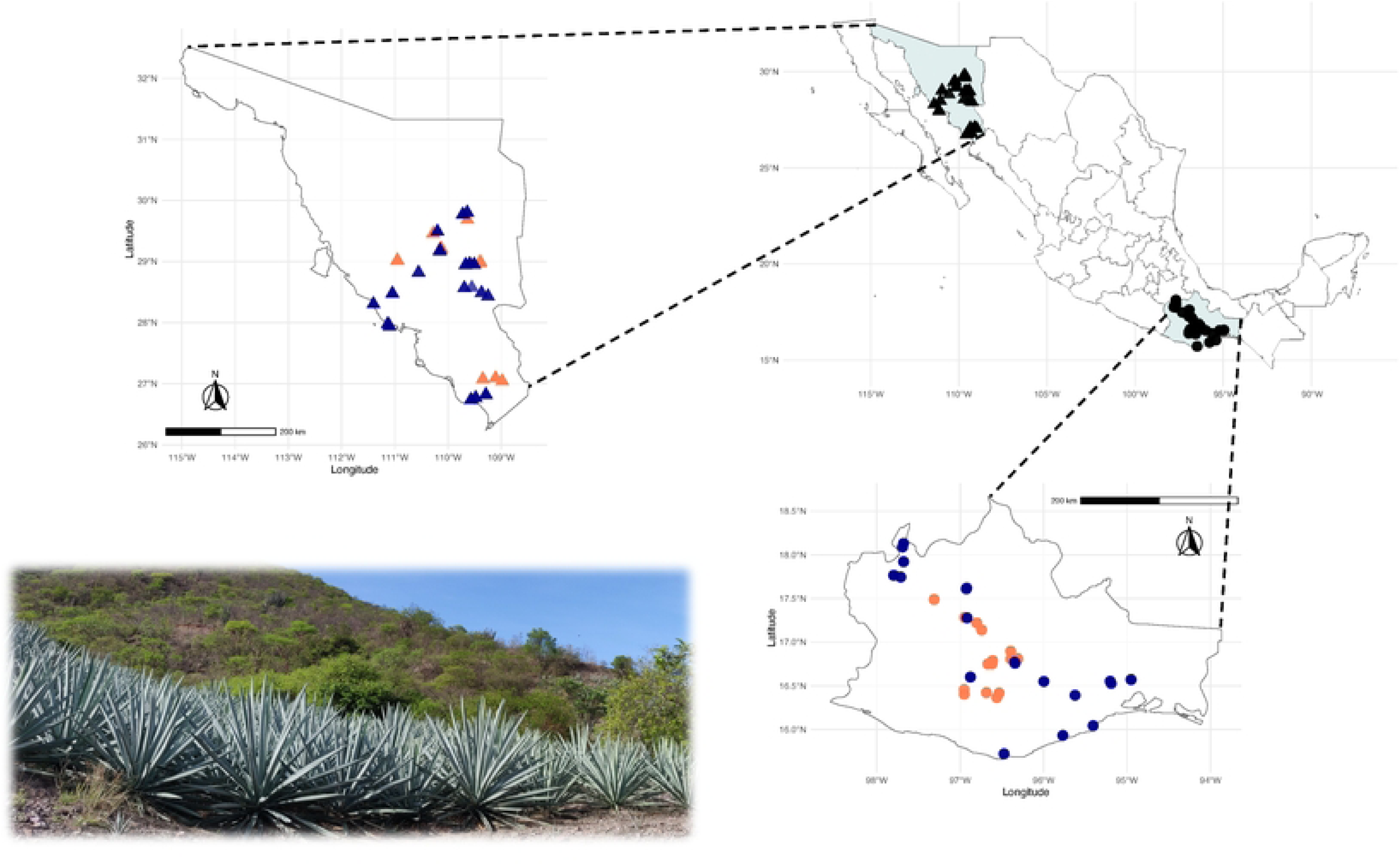
Map of the wild and cultivated *Agave angustifolia* collected in the states of Sonora and Oaxaca. The localities are colored as cultivated (dark orange) and wild (dark blue). In the left corner is a picture of cultivated Espadín collected in the Oaxaca state. In the right corner is a map of Mexico, with the Sonora and Oaxaca states marked in blue, showing all sampled localities in black, using triangles for Sonora and circles for Oaxaca.

The downloaded samples corresponded to GBS (Genotyping by Sequencing, [61]) data. In all the samples, DNA was digested with the two restriction enzymes (PstI/MspI) and sequenced using 1 × 150 Illumina NovaSeq at the University of Wisconsin Gene Core. For this new study, reads were mapped to a recently published genome of an *Agave* [62] using bwa-mem2 version 2.2.1 [63,64]. Mapping output files were converted to BAM and sorted with samtools version 1.19 [65]. Single nucleotide polymorphisms (SNPs) were first called for each individual using BCFtools [66]. Raw SNP filtering was performed using VCFtools version 0.1.16 [67] and plink version 1.90b6 [68]. At this step, only biallelic loci with a minimum mean depth of 10 and a maximum mean depth of 200 were kept. Additionally, we set a minor allele count at eight and excluded sites based on the proportion of missing data, keeping sites with no more than 10% missing data (--max-missing 0.9). We then removed variants significantly deviating from HWE (p ≤ 0.05 after Bonferroni correction) and sites in high Linkage Disequilibrium (r^2^>0.2 within 50 bp).

### Genomic diversity and relatedness

Although diversity estimation for clonal crops may not be straightforward [60], we employed several diversity metrics to find out how the management and domestication process have changed the cultivated plants and how habitat loss and extraction of wild seeds and plants may have affected the wild populations of *A. angustifolia*. We estimated observed and expected heterozygosity for each group. For each individual, we estimated multilocus heterozygosity and the Fhat3 inbreeding index using the R package InbreedR [69] and plink. We calculated per-site nucleotide diversity among samples of each management type and each state separately using the “--site-pi” function in VCFtools.

To evaluate how linkage disequilibrium (LD) decays with physical distance along the genome in wild and cultivated *A. angustifolia*, we calculated *r*^2^ values among all pairs of loci located within the same chromosome, separately for each group, using plink. LD decay was then visualized by binning the distance among SNPs into 1 kb intervals and plotting *r*^2^ values for each bin using R [70]. We also estimated the relatedness among each pair of individuals within each wild and cultivated group. The relatedness was estimated using a function within vcftools -- relatendess2 [71].

### Genetic differentiation

To evaluate the degree of genetic differentiation between wild and cultivated *A. angustifolia*, we calculated pairwise *F*_ST_ values using the R package stampp [72]. We also used the R package SNPrelate [73] to explore genomic divergence among individuals using principal component analysis (PCA). We constructed a distance tree with the R package poppr [74] to visualize the genetic relationships among samples and determine the number of genetic groups. The distance tree was based on the NJ algorithm, with 1000 bootstrap replicates to assess branch support.

We also used fastStructure v 1.0 [75], which was run with default settings, and 100-fold cross-validation on the 207 samples, testing for the best number of groups of populations (K), ranging from K = 1 to 10 with five repetitions for each K value. The best K value was chosen based on the highest marginal likelihood. The mean population membership probability was manually plotted in R based on the Q-matrix values produced by fastStructure. Finally, we compared allele frequency distribution between wild and cultivated individuals using vcftools and R.

### Effects of domestication

To evaluate the strength of a potential population bottleneck during *A. angustifolia* domestication and management, we identified alleles that were (1) private (unique) to the wild population, (2) shared between the wild and domesticated plants, or (3) private to each domesticated population (novel or increased in frequency). The allele frequency threshold to consider an allele present was set at 0.01. To determine the identities of polymorphisms private to the domesticated populations, the 150bp sequences contigs containing private SNPs were compared to a custom database using the BLASTx search tool with a minimum E value of 1 × e−10 [76]. The custom database included all the plant protein sequences available at UniProt (https://www.uniprot.org) and was constructed with makeblastdb [77]. The BLAST results were evaluated to determine each gene’s identity using the UniProt ID mapping tool (https://www.uniprot.org/id-mapping, accessed in January 2025).

Furthermore, we used three complementary approaches to search for outlier loci between wild and cultivated plants [78]. We used the R package outflank [79], which identifies *F*_ST_ outliers or loci with atypical values of *F*_ST_ by inferring a distribution of neutral *F*_ST_ using likelihood on a trimmed distribution of *F*_ST_ values. The outflank analysis was performed separately for Espadín and Bacanora samples. We created two data sets, one containing all wild and Espadín samples and another one containing all wild and cultivated Sonoran samples. In this analysis, the ‘number_of_samples’ parameter was set to three, and then to 53 in the case of Espadín and 51 in the case of Sonoran cultivated agaves (i.e., a number equal to the genetic groups identified with PCA, and the numbers of sampled sites). The ‘LeftTrimFraction’ and ‘RightTrimFraction’ were left at the default setting. The Hmin parameter was set at 0.05. The false discovery rate threshold for calculating q-values was set at 0.01 [79].

We also used the Bayesian method implemented in BAYESCAN V2.1 [80]. BAYESCAN tests whether subpopulation-specific allele frequencies, measured as F_ST_, are significantly different from the allele frequency within the shared gene pool and gives a posterior probability (alpha) to a model in which selection explains a difference in allele frequencies better than a null model. BAYESCAN was run using prior odds set at 100 [81]. A false discovery rate (FDR) of 0.05 was used, with the caveat that although this reduces the number of false positives, true selection signals may be missed [80].

Finally, we used LFMM [82] to identify outlier loci using a Bayesian mixed model with management type and population variables included as fixed effects. We built the model using the lfmm_ridge function of the R package LFMM [83]. We explored the number of latent factors (K value) from 2 to 4 and used regularization parameters (lambda) set to 1e-5 to minimize predictor error estimated by a cross-validation method. P-values were calibrated using the genomic control method, and the false discovery rate (q-value) was calculated following the Benjamini–Hochberg procedure in the qvalue R package [84]. Associations between an SNP and management with a q-value < 0.05 were considered statistically supported.

## Results

After reads alignment and SNP calling against the reference agave genome, a total of 3,938,407 variants were called. After filtering with VCFtools, our final data set consisted of 207 *A. angustifolia* samples (S1 Table) genotyped with 51,560 SNPs. The data included 47 individuals of cultivated *A.angustifolia* from Oaxaca (Espadín), 42 samples of cultivated *A. angustifolia* from Sonora (Bacanora agave), and 56 and 62 wild samples from Sonora and Oaxaca states, respectively. The average sequencing depth per individual was 25.3, and the average missingness at the individual level was 1.75%.

### Genomic diversity

The mean observed (*Ho*) and expected heterozygosity (*He*) of the analyzed groups of plants are shown in Table 1. The lowest *Ho* of 0.12 (SD: 0.12) was observed in wild samples from Oaxaca, whereas a slightly but significantly (after a Bonferroni correction) higher value (*Ho*=0.13, SD: 0.12) was found in wild samples collected in Sonora. The highest values of expected heterozygosity (*He*=0.15, SD: 0.13) were estimated for wild samples from Sonora, while the diversity was less than half of it in the cultivated samples from Oaxaca (*He*=0.07, SD: 0.15).

**Table 1.** Diversity estimates with corresponding standard deviation values (SD) for wild and cultivated *Agave angustifolia* in the Oaxaca and Sonora states estimated using 51,560 SNPs.

| | $N$ | $\pi$ | $H_o$ | $H_e$ | Fhat3 | MLH | Mean $r^2$ | Relatedness |
| --- | --- | --- | --- | --- | --- | --- | --- | --- |
| *A. angustifolia*<br>wild Sonora | 56 | 0.15 (0.13) | 0.13 (0.12) | 0.15 (0.13) | 0.13 (0.05) | 0.13 (0.009) | 0.02 (0.05) | 0.01 (0.07) |
| *A. angustifolia*<br>wild Oaxaca | 62 | 0.13 (0.14) | 0.12 (0.12) | 0.13 (0.14) | 0.15 (0.04) | 0.12 (0.007) | 0.02 (0.05) | 0.01 (0.07) |
| *A. angustifolia*<br>cultivated Oaxaca | 42 | 0.07 (0.16) | 0.12 (0.29) | 0.07 (0.15) | -0.06 (0.04) | 0.12 (0.01) | 0.2 (0.29) | 0.41 (0.10) |
| *A. angustifolia*<br>cultivated Sonora | 42 | 0.15 (0.13) | 0.13 (0.13) | 0.15 (0.13) | 0.14 (0.04) | 0.13 (0.003) | 0.03 (0.07) | 0.02 (0.09) |

We observed a slight heterozygote deficiency in wild samples of both states and in the cultivated Sonoran plants, whereas a heterozygote excess was only observed in cultivated Espadín (Table 1). We found that wild agaves of both states and cultivated plants from Sonora were moderately inbred according to the Fhat3 inbreeding index, with similar *f* averaging 0.13 (SD: 0.05) for wild plants from Sonora, 0.14 (SD: 0.04) for cultivated agaves from Sonora, and *f* averaging 0.15 (SD: 0.04) for wild plants from Oaxaca. The highest inbreeding values (0.25 and 0.24) were found in one wild plant from Sonora and one cultivated individual from Sonora. In wild individuals from Oaxaca, *f* was negative, -0.06 (SD: 0.03).

Individual-based multilocus heterozygosity (MLH) was similar between wild and cultivated plants within each state but slightly higher for Sonoran populations (Table 1). Cultivated Espadín had considerably lower average nucleotide diversity (0.07, SD: 0.15) compared to wild samples from Oaxaca (0.13, SD: 0.13) and wild samples from Sonora (0.15, SD: 0.13). Meanwhile, Sonora’s managed samples had the same nucleotide diversity levels as wild plants from the region.

### Linkage disequilibrium and relatedness

Samples were divided into four groups (cultivated and wild plants belonging to two states), and pairwise LD was estimated within the gene pool of each of these groups. The strength of LD was considerably different between clusters, as reflected by the mean *r^2^* values of 0.20 (SD: 0.29) and 0.02 (SD: 0.04) obtained for the cultivated Espadín and the wild plants from Oaxaca, respectively.

Moreover, LD did not decay with increased distance between markers in the cultivated plants from Oaxaca. In contrast, LD rapidly decayed and maintained relatively low levels in the wild samples. The LD decay was similar between wild samples from the two states (Fig 2). The patterns of LD decay in cultivated plants from the Sonora state were similar to those found in wild plants; however, the mean LD was higher at 0.034 (SD:0.07) and did not decay to the level observed in wild plants (Fig 2).

**Fig 2.**
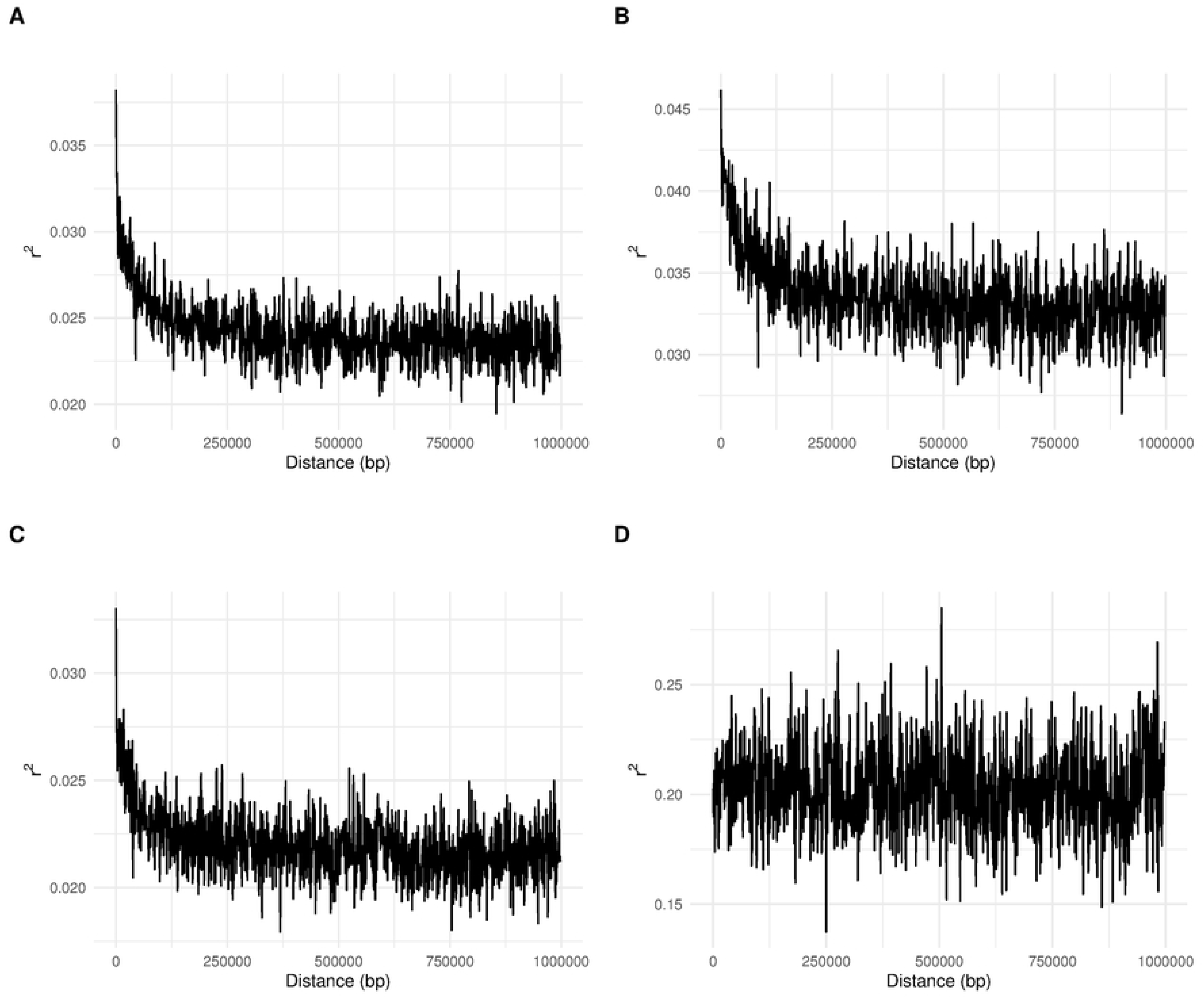
Scatter plot of LD decay (r^2^) against the distance in base pairs for pairs SNP for (**A**) wild Sonoran populations, (**B**) managed Sonoran populations, (**C**) wild populations from Oaxaca, and (**D**) managed Espadín from Oaxaca.

The relatedness estimates also differed drastically between the analyzed group of samples (Fig 3), suggesting different forms of propagation for wild and cultivated individuals. The highest relatedness was found among Espadín samples (mean 0.41, SD 0.1), suggesting that almost all the samples may be considered unique clone with very little genetic differentiation among them. On the other hand, cultivated samples from the Sonora state presented low relatedness (mean 0.02, SD 0.087), which was very similar to the results for wild samples (Fig 3). Nevertheless, few samples, generally collected at the same plantation, presented high relatedness among them (Fig 3). The mean relatedness among wild samples from Oaxaca was low (mean 0.01, SD 0.067), suggesting that breeding between related individuals is uncommon. Similar results were found for the wild samples from the Sonora state, with a mean relatedness of 0.01, SD 0.073 (Fig 3). However, there was a variation in relatedness estimate among samples, with some presenting high levels of genetic affinity, particularly samples from the same location.

**Fig 3.**
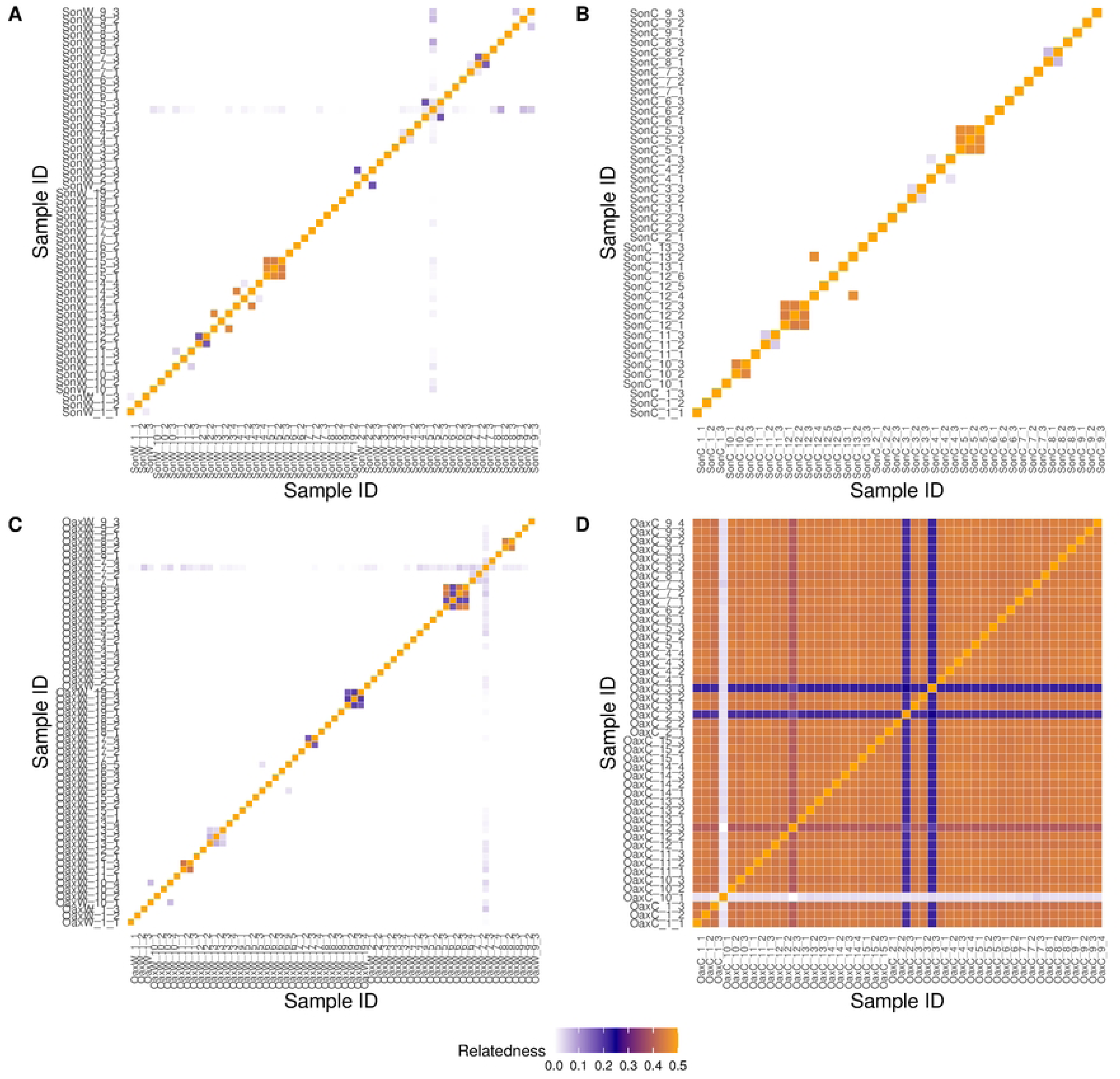
The diagram of relatedness coefficient as estimated within vcftools for (A) 56 wild and (B) 42 managed *A. angustifolia* individuals from the state of Sonora, (C) 62 wild, and (D) 47 managed *A. angustifolia* individuals from the state of Oaxaca.

### Genetic differentiation

As with diversity levels, genomic differentiation differed for Oaxaca and Sonoran plants. In the case of *A. angustifolia* in Sonora, no genomic differentiation was found between wild and cultivated individuals (Fig 4), and the *F*_ST_ was significant but low (*F*_ST_=0.005, 95% CI 0.0049-0.0054). On the PCA and NJ tree, the cultivated samples from Sonora were dispersed within wild samples, suggesting that producers from different parts of Sonora still incorporate local wild plants and seeds into cultivation. The Espadín used for producing mezcal in Oaxaca presented significant levels of differentiation from wild samples from Oaxaca and Sonora (*F*_ST_ =0.095 – 0.106).

**Fig 4.**
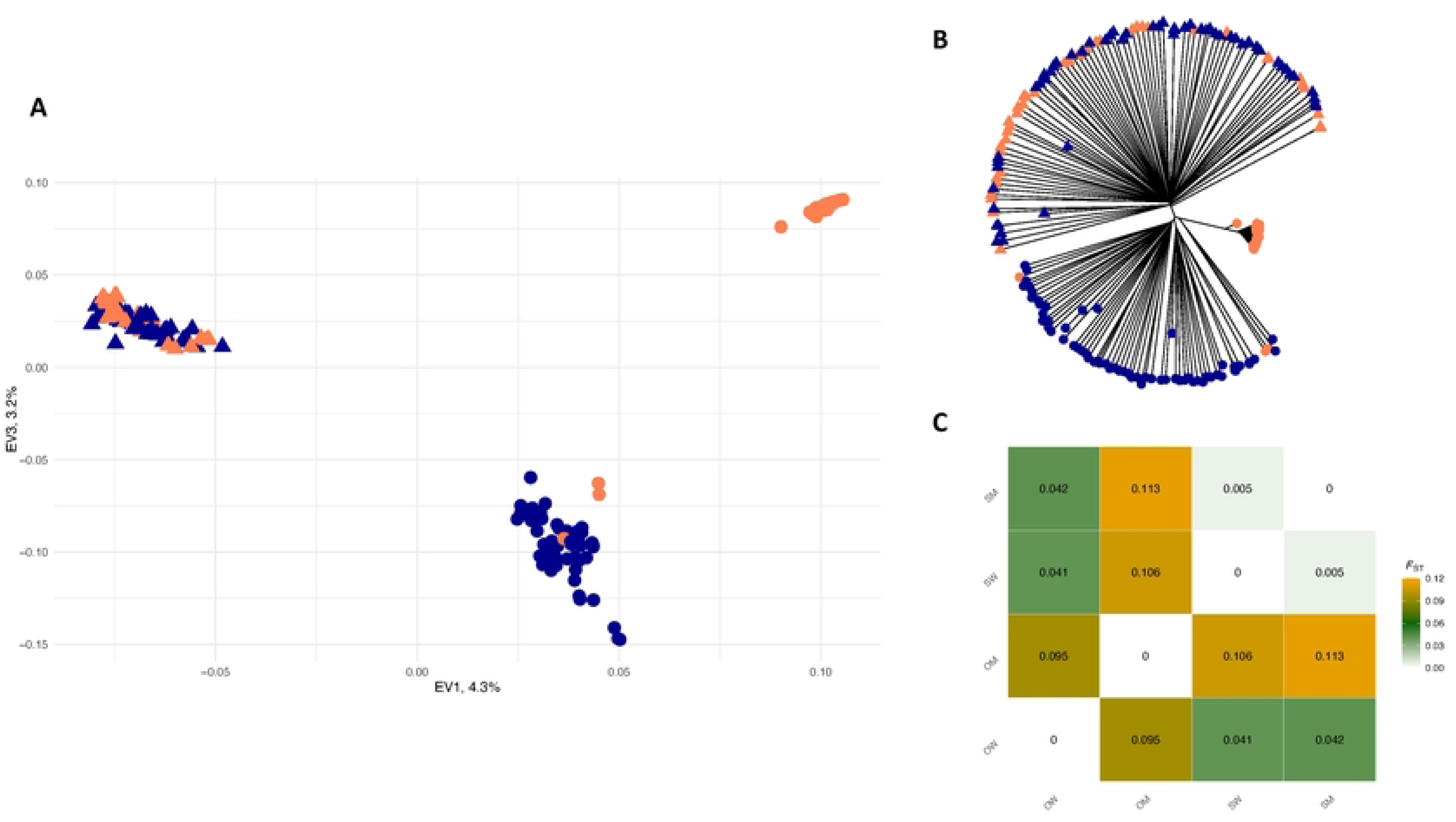
Population genetic structure of wild and cultivated *A. angustifolia* from Sonora and Oaxaca states, Mexico, based on 51,560 genome-wide SNPs. (A) Principal component analysis (PCA) of the individuals of *A. angustifolia*. (B) Neighbor-joining (NJ) network of 207 individuals of *A. angustifolia*. PCA and NJ tree tips are colored according to the management type of the samples: dark blue for wild and dark orange for managed. Shapes on PCA and NJ tree tips correspond to the state where samples were collected: circles to Oaxaca and triangles to Sonora. (C) Pairwise *F*_ST_ differences among wild and cultivated samples of *A. angustifolia* from Sonora and Oaxaca states, Mexico. Colors represent *F*_ST_ values from the lowest of 0 in white to the highest of 0.12 in orange. Abbreviations: OW-Oaxaca Wild, OM – Oaxaca Managed, SW - Sonora Wild, and SM - Sonora managed.

Espadín was clearly differentiated according to the PCA (Fig 4), forming a group at the upper right corner, separated from wild samples of Oaxaca and wild and cultivated samples of Sonora (Fig 4A). However, we must point out that the two main PCA axes explained a relatively low (a total of 7.5%) percentage of the variation. The NJ tree revealed three samples designated as Espadín clustered along wild samples of Oaxaca. In contrast, the remaining Espadín samples formed a tight cluster of samples on a separate branch. Moreover, we found a significant divergence between wild samples from Sonora (triangles in the figure) and Oaxaca (circles in the figure).

All the above analyses confirmed that wild samples from Sonora are genetically distinct from those collected in Oaxaca, with a significant *F*_ST_ of 0.041 (95% CI 0.04-0.041) between them and no admixed individuals.

The Bayesian clustering algorithm of fastStructure pointed to K = 3 as the number of groups that maximized the log-marginal likelihood lower bound. These results were identical to PCA, where we identified three main genetic clusters (S1 Fig). The three groups were clearly separated with basically no or very little admixture and corresponded to (i) wild and cultivated samples from Sonora, (ii) wild samples from Oaxaca, and (iii) cultivated samples from Oaxaca (S1 Fig).

The genomic differentiation between wild and cultivated Espadín samples was also visible in changes of correlation between allele frequencies (Fig 5A and C). Many alleles presented in high frequency in wild samples were presented in low frequency in cultivated Espadín and vice versa (Fig 5C). On the other hand, the correlation was almost perfect in the cultivated Sonoran samples, with similar changes in allele frequencies between the two groups of samples (Fig 5A).

**Fig 5.**
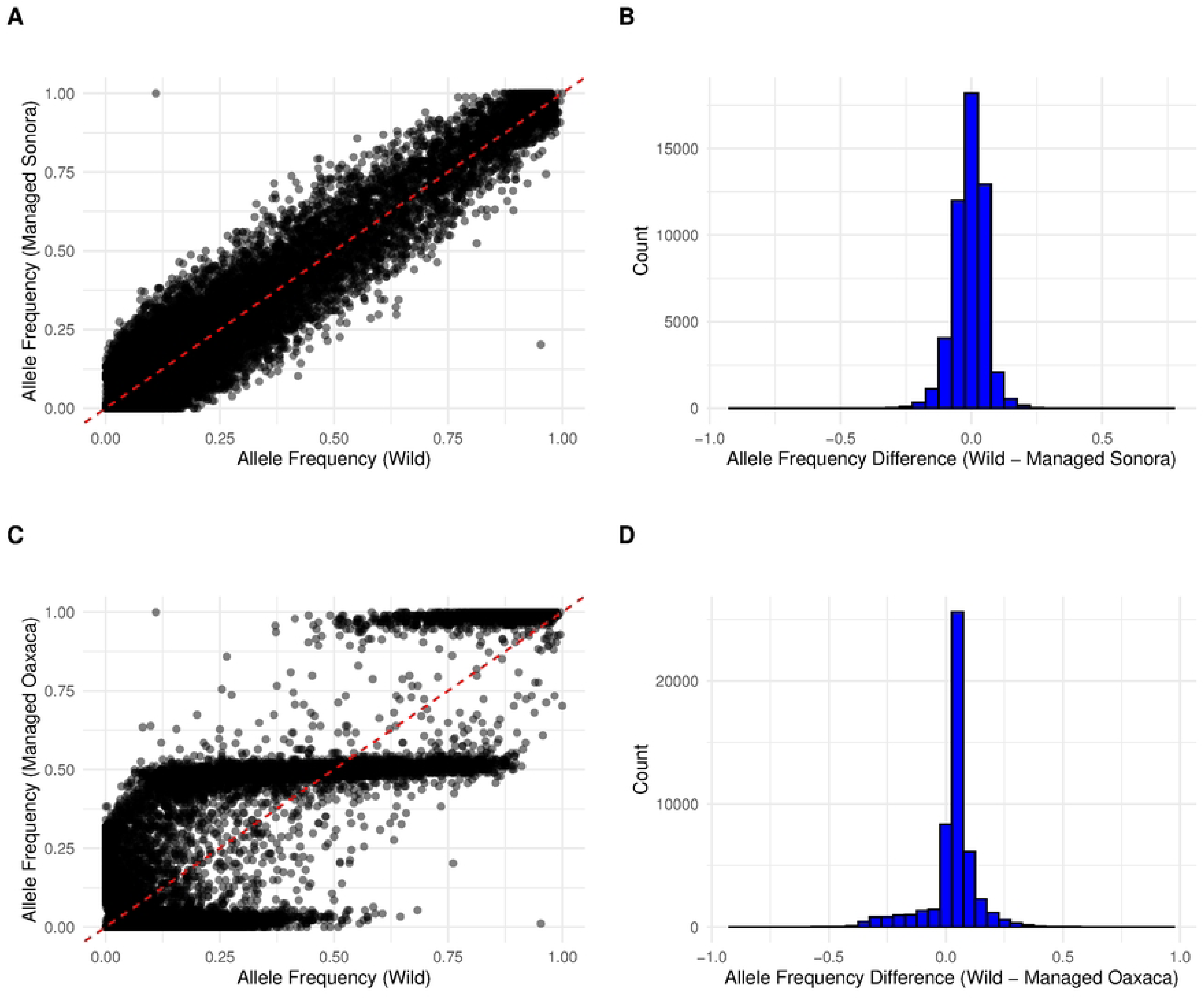
Comparison of the allele frequencies (A) and (B) the distribution of allele frequency differences between wild and managed *A. angustifolia* from the Sonora state and (C) and (D) between wild and managed *A. angustifolia* from the Oaxaca state.

### Effects of domestication

The allele frequency analyses revealed that in the case of Espadín, the number of alleles private to the wild population (i.e., alleles lost during domestication) exceeds the number of alleles shared (retained after domestication), or private to the domesticated population (putative novel alleles since domestication, or alleles that increased in frequency) (Fig 6 B).

**Fig 6.**
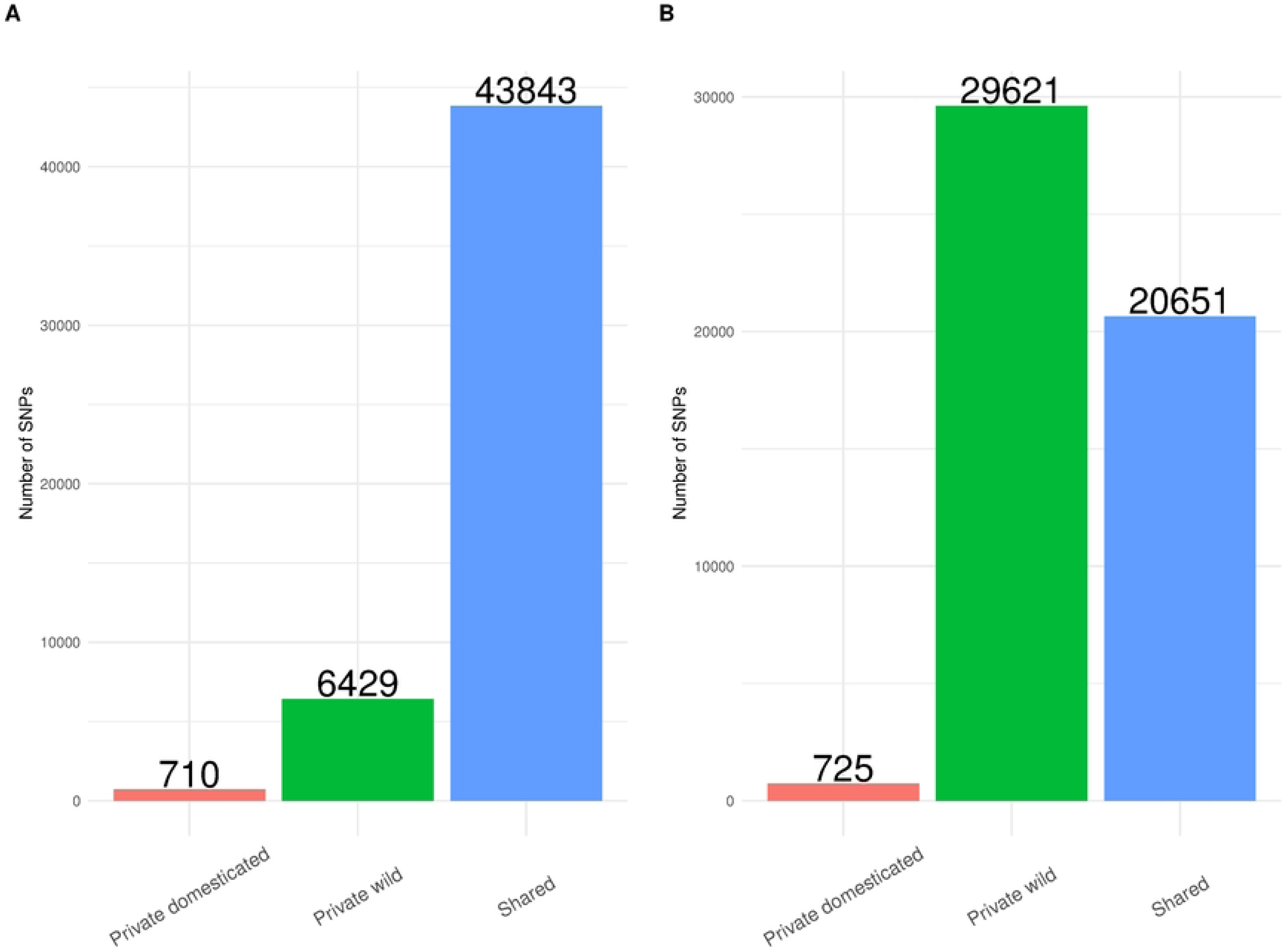
The number of alleles identified as shared, private to domesticated populations, and private to the wild samples for managed (A) *A. angustifolia* in Sonora and (B) managed Espadín in Oaxaca.

In the cultivated Bacanora samples, the number of alleles private to the wild population (i.e., lost during domestication) was much lower than the number of alleles shared (retained during domestication). Overall, we estimated that approximately 87.2% of wild alleles were retained in cultivated Sonoran samples, and 41% of wild alleles were retained in cultivated Espadín from Oaxaca (Fig 6). We found that a relatively small proportion of variation was unique to domesticated lineages, 1.59% in the case of the Sonora sample and 3.5% in the case of cultivated Espadín (Fig 6).

We also found shared alleles between cultivated samples; most unique alleles were found in cultivated Sonoran samples (26,456), 3,297 alleles were unique to Espadín, whereas 18,097 were shared between both cultivated plants.

When the BLAST sequences belonging to the alleles that were unique to cultivated Espadín and Bacanora agaves were compared against the UniProt database, we recovered 118 BLAST hits for Espadín and 146 hits for Sonoran samples (S1 and S2 Tables). We removed uncharacterized proteins and repetitive hits, reducing the number of successful hits to 85 for Espadín and 105 in Sonoran samples (S1 and S2 Tables). Some of the alleles unique to cultivated Espadín belong to proteins and transcription factors that have been reported as important for plant growth, seed development, defense, and tolerance to stress [85,86,87,88,89].

On the other hand, no outliers were detected for the managed populations, independent of the three methodologies used.

## Discussion

Domestication has transformed wild plants to the extent that some crop species are phenotypically very different from their ancestors [8,90]. Although agaves have served people for multiple purposes for thousands of years, the plants generally have not changed much. This lack of apparent morphological change may be attributed to a combination of factors, such as the relatively low strength of artificial selection, the long generation time of agave species, high phenotypic plasticity, and the possibility of asexual reproduction [25,91,92]. However, currently, the tequila and mezcal industries are pushing artificial selection, seeking to increase size and sugar content, and shorten generation time. To achieve these objectives, one of the primary practices adopted by agave producers is preventing blooming and widespread reliance on clonal propagation with a selection of the best clones [58,59,60].

Based on our findings, we hypothesize that Espadín recently aroused through farmer selection of a clonal lineage with desirable mezcal production attributes (i.e., related to production and hybrid vigor) from a cross between genetically distinct wild populations or by hybridization between wild and cultivated variety of *A. angustifolia* that are still common in Jalisco [45,93]. This process is sometimes called “instantaneous domestication” [94,95]. The following evidence seems to support this hypothesis. First, we found that Espadín samples were represented by a single heterozygous genotype (except for three individuals that were classified by PCA and NJ tree as wild) with extreme genetic homogeneity among samples and high linkage among loci. Second, we observed a considerable genetic differentiation between managed Espadín and wild populations from both states, which is unexpected, given the alleged recent origin of Espadín from local wild plants. The origin from ancestral recombining populations would have generated conflicting topologies [95] and cultivated samples distributed along several tree branches of wild samples, which was not observed in the case of Espadín. Finally, although we found alleles that increased in frequency in cultivated Espadín, none of the outlier analyses detected loci under selection.

On the other hand, the genomic composition of cultivated Bacanora agave was apparently highly influenced by the region’s social and political history. We suggest that the ban on cultivation and distillation that lasted over 70 years, along with the relatively lower popularity of the beverage compared to tequila and other mezcals, allowed cultivated Bacanora to maintain the genetic diversity found in wild populations of the regions. Additionally, our findings indicated that farmers even now include local wild seeds and plants in cultivation.

### Genomic diversity

The management practices in agave can vary significantly, ranging from low-scale agroforestry to extensive monoculture fields that use cloned plants [96,97,98]. This is particularly evident in *A. tequilana* var. *azul* in Jalisco and Guanajuato and Espadín in Oaxaca. Such agricultural systems are highly intensified and are characterized by large areas planted with a single species, disruption of the natural growth cycle of the plants (including the prevention of flowering and pollination), and the use of agrochemicals [29,99,100]. It has been suggested that the time under cultivation and the intensity of management affect the speed and amount of genetic diversity lost in agaves [101,102]. Agave species with relatively recent or low-intensity management, such as *A. marmorata*, *A. potatorum*, *A. cupreata, A. americana*, *A*. *salmiana, A. inaequidens* and *A*. *mapisaga* contain the same or similar levels of diversity as their wild conspecifics [92, 102,103,104,105]. Others, such as *A. tequilana* var. *azul*, have lost much of their genetic diversity. For instance, [101] found that all 23 analyzed *A. tequilana* var. *azul* plants shared the same genotype. Interestingly, the average levels of genetic variation found in other traditional tequila cultivars are similar to those observed in wild populations of *A. angustifolia* [93,101].

We found contrasting levels of genomic diversity in cultivated Espadín and Bacanora agaves (Table 1). Espadín had high observed heterozygosity and a negative inbreeding index, but it had the lowest expected heterozygosity, less than half of the values found in wild plants and cultivated Sonoran samples. Espadín also had the lowest nucleotide diversity values: half or even lower than those in other studied groups (Table 1). In addition, nearly all analyzed Espadín individuals from 15 different plantations could be regarded as a single heterozygous clone. The diversity found in Espadín is similar to that reported for *A. tequilana* var. *azul* and some landraces from Jalisco [45,101] and may be explained by the exponential demand for mezcal production of plants in the shortest possible time, following the tequila production model of cultivation practices promoting the use of clones.

In contrast, the genomic diversity of the cultivated plants in the Sonora region closely resembled that of their wild relatives, which is characteristic of agaves managed with low-intensity practices (Table 1). Thus, we did not detect any evident reduction in diversity in cultivated Sonora plants. These findings may be explained by a long-term ban on plantations in Sonora state, the relatively recent increase in Bacanora production, the lesser demand for the beverage on the international market, and the producer’s habit of including wild plants and seeds in cultivation.

However, we found a slight increase in relatedness estimates and the linkage disequilibrium among loci, which may indicate that Sonoran farmers are beginning to adopt management practices characteristic of *A. tequilana* var. *azul* and Espadín.

We, therefore, can conclude that the social and political histories of the Sonora and Oaxaca states have had an important effect on cultivated agaves’ genomic diversity. Moreover, given the evidence of reduced diversity and increased relatedness for cultivated Espadín and data on genetic diversity in *A. tequilana* var. *azul,* it is of utmost importance for Bacanora producers to avoid the fate of tequila and Espadín plants and maintain levels of genetic diversity that will allow to create new varieties, capable of coupling with changing climate and evolving pests.

Wild samples from the Sonora state had slightly higher levels of genomic diversity than wild plants collected in Oaxaca. For example, heterozygosity and nucleotide diversity were significantly lower in wild Oaxaca populations. However, the inbreeding index and relatedness were similar between wild groups and did not display significant differences. It is possible that the wild plants in Oaxaca are experiencing greater anthropogenic pressure or that the populations were always smaller than the Sonoran populations. However, more *A. angustifolia* populations will need to be analyzed to determine the base level genetic diversity for this species and explain how natural or human-induced forces may have affected patterns of genomic diversity distribution found in Sonoran and Oaxaca wild plants.

### Clonality and domestication bottleneck

The high levels of relatedness among Espadín individuals and high linkage disequilibrium among loci are the consequence of clonal propagation and the prevention of flowering, pollination, and genetic exchange among plants [106,107]. However, from an ecological and evolutionary perspective, asexual propagation is considered an adaptation and should not be viewed negatively. Asexual or clonal propagation is widespread in plants and offers several ecological benefits compared to sexual reproduction [108,109]. These advantages include the ability of clones to “search” for light, nutrients, and water, lower likelihood of death of clonal genotypes because the risk of mortality is spread among numerous shoots, and the ability of clonal propagules to disperse to new environments where, being larger than seeds and lacking dormancy, they can multiply rapidly and compete effectively with other species [108,109].

Besides the ecological advantage, clonal reproduction represents several benefits for farmers and has, therefore, been extremely important in the domestication of many crop species, including agave [106]. The most obvious advantage is that clonal propagation is the easiest way to multiply desirable genotypes. Clonal propagation ensures faster initial growth and a higher survival rate than seed propagation. In addition, clonal reproduction allows the instantaneous fixation of genotypes with agronomic value, ensuring that favorable genotypes are faithfully transmitted to the next generation of the crop. Clonal propagation can also help preserve highly heterozygous genotypes that show hybrid vigor. Individuals with advantageous mutations can also be easily identified in the field and rapidly propagated. Finally, clonal propagation helps to avoid gene flow between crops and their wild relatives, a gene flow that may increase the proportion of undesirable variants according to the producer’s perspective [106,110].

However, clonal propagation has several important disadvantages, such as the accumulation of deleterious mutations and the possible spread of systemic pathogens. Moreover, since recombination is absent during clonal reproduction, almost no new genetic variation is generated [106]. Moreover, a lack of sexual reproduction can limit crop yield and its potential for improvement [106]. For instance, in cereals such as rice or maize, rounds of self-pollination have been used to create pure inbred lines that, when crossed, produce uniform and vigorous hybrid seeds and increase productivity. However, similar progress is hard to achieve in clonally reproduced crops [108, 111].

In the cultivated Espadín, the negative consequences of the widespread adoption of clonal propagation are already noticeable, including a considerable loss of genomic diversity and extreme morphological and genetic homogeneity. We also found a significant divergence between wild and cultivated plants, with clear changes in allele frequencies (e.g., increasing from low-frequency alleles in wild plants to high frequency in cultivated individuals, or even the appearance of alleles not present or present in very low frequencies in wild plants). Indeed, we found several alleles private to Espadín or alleles that increased in frequency within the protein-coding genes responsible for important ecological processes that may be under artificial selection (S1 Table). We believe this means that the observed divergence is not simply a by-product of genetic drift but also a consequence of human selection and may involve adaptive divergence.

Supporting these ideas, we detected changes in allele frequencies, including in genomic regions known to be involved in several biological processes that may be under selection, for instance, in growth, stress tolerance, and immunological responses. Among them, we found the AT-hook motif nuclear localization (AHL), bHLH transcription factors, pentatricopeptide repeat proteins, WRKY proteins that play roles in plant growth and development, response, and adaptation to ecological stresses through protein-DNA and protein-protein interactions [87,88,112,113]

We also detected changes in allele frequency in regions responsible for plant defense, including chitinase enzyme production. Chitinases are defense enzymes expressed in plants in response to fungal pathogens [114]. LysM proteins that play a role in maintaining plant-microbe symbioses, such as the interaction between rhizobacterium and leguminous plants or the interaction between different plant roots with mycorrhizal fungi. These proteins can differentiate between beneficial and pathogenic microbes due to their ability to recognize molecular patterns of different pathogens [115]. Another interesting protein is Cytochrome P450 (CYP) 81, which is involved in herbicide metabolism and other processes in various cultivated plants, including rice, soybeans, and grapes [116].

However, the observed changes in allele frequency may also be explained by the alleles inherited from an ancestor (still unknown) of the cultivated Espadín. At this moment, we can only speculate regarding the real biological impact of these allelic changes. Future research in *A. angustifolia* and Espadín should focus on the adaptive divergence and negative consequences of clonal reproduction, including evaluating the levels of deleterious mutations, genetic load, and changes in pathogen burden.

### Genomic differentiation

We found clear genomic differentiation in cultivated Espadín and Bacanora agaves. According to our data and what the local people told us, the most important source of Bacanora plant germplasm in Sonora is the plants and seeds obtained by farmers in the vicinity. Subsequently, the collected plants are propagated by offsets (asexual reproduction), and seeds are germinated. The constant recruitment and propagation of local wild plants resulted in low differentiation between wild and cultivated individuals and a high number of shared alleles. In fact, the genomic differentiation for the Sonoran wild and cultivated plants is the same or even lower than those found in other cultivated agaves used for mezcal production, such as *A. potatorum* and *A. marmorata* [104,105].

This is related to the relatively recent increase in production, lower production level, lesser commercialization on the international level, lower involvement of big multinational companies in Bacanora cultivation, less influence of the tequila industry, and the historic habit of using wild plants and seeds [59]. For instance, Sonora’s annual production levels of 300,000 liters do not even come close to the mezcal production scale in Oaxaca, which reached 6,230,378 in 2022 and, although it dropped in 2023 to 3,798,753, is over 10 times higher than the production of Bacanora [44,117].

The longer time under cultivation and high-scale industrial use of Espadín in monoculture cannot completely explain its relatively high levels of genomic differentiation compared to the wild plants from Oaxaca. For example, [45,101] could not find genetic evidence to delimitate cultivars of *A. tequilana*, *A. rhodacantha*, or wild *A. angustifolia* collected in Jalisco state. However, these studies did not report *F*_ST_ values between *A. tequilana* var. *azul* and wild and cultivated agaves, so it is difficult to compare our results in the framework of these studies.

Given the relatively high genetic differentiation found in Espadín, its divergence from wild samples on the PCA and NJ tree (Fig 4), and differences in allele frequencies compared to wild samples (Fig 5), there are several possible explanations for the origin of these plants. Espadín may have originated from the crossing between two genetically distinct wild populations. However, these populations have not been sampled in our analysis or have disappeared due to habitat loss. Indeed, the extinction of some wild populations in Oaxaca is possible as, in the last decades, the state has suffered major anthropogenic disturbances, including extensive deforestation and natural habitat conversion to agriculture, more recently to plant mezcal agaves [118,119]. Another possibility is that Espadín originated from a cross between genetically distinct wild populations of *A. angustifolia* from very different geographic regions. However, given that our analyzed sampling is restricted to only two states, we cannot explore this hypothesis further.

To elucidate Espadín’s genomic origin, future studies should sample wild *A. angustifolia* throughout its distribution range and determine which wild samples may have given rise to it. Now, we can only conclude that, given the genetic differentiation, wild samples from Sonora or Oaxaca do not seem to be a valid candidate as Espadín’s ancestor. A plausible hypothesis is that Espadín originated as a hybrid between wild plants and a cultivated *A. angustifolia* variety or between two cultivated *A. angustifolia* varieties.

However, genomic analyses of *A. angustifolia* cultivars will be needed to explore this hypothesis, which is beyond the scope of the present work. At this moment, we can conclude that the origin of cultivated Espadín was a complex process that may have included cross or crosses between wild plants from different geographic regions or alternatively with already developed cultivars.

### Future management and conservation

The ubiquity of clonal propagation in agriculture, in general, and in agave cultivation in particular, reflects its value to farmers in selecting and rapidly multiplying favorable genotypes. However, this strategy may become suboptimal in the near future. Mexico is already experiencing climate change, and climate effects are projected to become more intense throughout this century [120,121]. Thus, in the states of Oaxaca and Sonora, increases in temperature and drought intensity have already been documented [122,123,124].

Farmers will rely on three main natural-biological strategies for their crops to survive major shifts in biotic and abiotic conditions under changing climate: use available phenotypic plasticity; expect the crop to adapt to the new environments with its standing genetic variation; or hope the crop can get useful genetic variation via gene flow [125,126,127]. Indeed, many natural plants can adjust their phenotype in response to changes in environmental conditions; this kind of response does not require changes in gene frequencies (i.e., evolution). Such phenotypic shifts can allow current populations to maintain their fitness as conditions change. However, agaves will need to respond plastically to several environmental changes simultaneously. Unfortunately, a particular plastic response may be beneficial in response to one environmental cue but detrimental in response to another [126].

In agaves, studies on the reaction norm of landraces or clones under various conditions – across years, locations, or with experimental treatments mimicking climate change – are lacking but urgently needed, as they will help to understand the range of environments under which landrace populations may be able to maintain fitness.

Natural migration or gene flow can also facilitate adaptation and maintenance of productivity under climate change because it can introduce novel variation on which selection can act [125,128]. However, current agricultural practices in Espadín cultivation hinder flowering and pollination in all plantations, and therefore, according to our data and observations, apparently, no gene flow exists between wild and cultivated plants in Oaxaca.

In Sonora, on the other hand, farmers often include local plants and seeds in cultivation, with some producers permitting a portion of their plants to flower (some of these producers participate in specific conservation programs). These factors suggest that Bacanora agave may be better prepared to handle climate change.

Changes in environmental conditions could also drive crop landraces to evolve or ‘keep up’ with climate change by natural selection on beneficial traits, changing allele frequencies at the loci that control these traits at a population level. For natural selection to occur, there must be sufficient and useful standing genetic variation within the crop population to allow for adaptation to the new environmental conditions [126]. However, in clonally propagated crops, such as agave, more genetic variation is found among populations than within them. As a result, selection may eliminate entire populations (i.e., clones) rather than cause genetic shifts within populations. Unfortunately, current agronomical practices in tequila and mezcal cultivation rely heavily on a few clonally propagated landraces (i.e., *A. tequilana* var. *azul* and Espadín), suggesting that climate change may negatively affect the plantations, leading to catastrophic consequences.

Besides climate change, after more than 200 years of the agave-derived alcohol industry, several ecological challenges have intensified in recent years. Agave monocultures have altered pollination corridors, particularly for nectarivorous bats, which has promoted a widespread problem affecting several agave species in neighboring regions [129,130,131,132]. Furthermore, the reduction of genetic diversity and the loss of cultivars and local varieties threaten the evolutionary potential of different agave species [12,133]. Forest cover loss is also a significant problem; in some parts of Oaxaca, the number of agave plantations has significantly increased, replacing natural vegetation and traditional crop species [119,134]. Besides ecological challenges, the mezcal industry has an important social impact, which has not always been considered positive. For example, high demand for mezcal plants has led to conflicts over water [135]. Furthermore, industrial monopolies in the agave industry have led to diseases, soil erosion, chemical contamination, and shifts from traditional crops to agave, causing social injustice for small producers and indigenous peoples [34,45,136].

We suggest that under a climate change scenario characterized by warmer and drier conditions, the balance between clonal propagation and sexual recombination for Espadín should favor sexual reproduction. However, the lack of evolutionary perspectives among agave producers regarding clonal plant breeding will require collaboration between producers, who may still have a useful repository of agave germplasm, and researchers. Together, we believe we can work to identify or develop genotypes that are better suited for future conditions.

Finally, further research should focus on wild *A. angustifolia* to determine how it responded to major past ecological changes and how it will respond to ongoing climate change. It should also identify which populations carry alleles that will allow them to withstand and adapt and can be used in cultivated plant breeding.

## ACKNOWLEDGMENTS

We thank Rosalinda Tapia for specialized technical support in the laboratory and Guillermo Sanchez de la Vega, Karen Ruiz Mondragon, Miguel Rivera-Lugo, Francisco Molina Freaner and José Fulgencio Martínez Rodríguez, all from Universidad Nacional Autónoma de México (UNAM), for crucial help in field sampling. We are particularly thankful to all the maguey and mezcal producers and families in the municipalities visited. This study was analyzed and written during the sabbatical leave of LEE 2024-2025, and he wishes to thank the UNAM for allowing him to have this sabbatical.

## AUTHOR CONTRIBUTIONS

AK, JL, and EAP designed the study; AK analyzed the data; AK and JL wrote the manuscript; all authors approved the final version of the manuscript.

